# Causal gene regulatory network inference using enhancer activity as a causal anchor

**DOI:** 10.1101/311167

**Authors:** Deepti Vipin, Lingfei Wang, Guillaume Devailly, Tom Michoel, Anagha Joshi

## Abstract

**Motivation:** Transcription control plays a crucial role in establishing a unique gene expression signature for each of the hundreds of mammalian cell types. Though gene expression data has been widely used to infer the cellular regulatory networks, the methods mainly infer correlations rather than causality. We propose that a causal inference framework successfully used for eQTL data can be extended to infer causal regulatory networks using enhancers as causal anchors and enhancer RNA expression as a readout of enhancer activity.

**Results:** We developed statistical models and likelihood-ratio tests to infer causal gene regulatory networks using enhancer RNA (eRNA) expression information as a causal anchor and applied the framework to eRNA and transcript expression data from the FANTOM consortium. Predicted causal targets of transcription factors (TFs) in mouse embryonic stem cells, macrophages and erythroblastic leukemia overlapped significantly with experimentally validated targets from ChIP-seq and perturbation data. We further improved the model by taking into account that some TFs might act in a quantitative, dosage-dependent manner, whereas others might act predominantly in a binary on/off fashion. We predicted TF targets from concerted variation of eRNA and TF and target promoter expression levels within a single cell type as well as across multiple cell types. Importantly, TFs with high-confidence predictions were largely different between these two analyses, demonstrating that variability within a cell type is highly relevant for target prediction of cell type specific factors. Finally, we generated a compendium of high-confidence TF targets across diverse human cell and tissue types.

**Availability:** Methods have been implemented in the Findr software, available at https://github.com/lingfeiwang/findr

## 1 Introduction

Despite having the same DNA, gene expression is unique to each cell type in the human body. Cell type specific gene expression is controlled by short DNA sequences called enhancers, located distal to the transcription start site of a gene. Collaborative efforts such as the FANTOM (Andersson *et al*., 2014) and Roadmap Epigenomics (Kundaje *et al*., 2015) projects have now successfully built enhancer and promoter repertoires across hundreds of human cell types, with an estimated 1.4% of the human genome associated to putative promoters and about 13% to putative enhancers. Enhancers physically interact with promoters to activate gene expression. Although the general rules governing these interactions (if any) remain poorly understood, experimental techniques such as chromosome conformation capture (3C, 4C) combined with next generation sequencing (Hi-C) (Mifsud *et al*., 2015) as well as computational methods based on correlations between histone modifications or DNase I hypersensitivity at enhancers with the expression of nearby promoters (Thurman *et al*., 2012; He *et al*., 2014) are continually improving at predicting enhancer-promoter interactions. In contrast, understanding how the activation of one gene leads to the activation or repression of other genes, i.e. uncovering the structure of cell-type specific transcriptional regulatory networks, remains a major challenge. It is known that promoter expression levels of transcription factors (TFs) co-express and cluster together with promoters of functionally related genes (Forrest *et al*., 2014), but without any additional information such associations are merely correlative and do not indicate a causal regulation by the TF.

Statistical causal inference aims to predict causal models where the manipulation of one variable (e.g. expression of gene A) alters the distribution of the other (e.g. expression of gene B), but not necessarily vice versa (Pearl, 2009). A key role in causal inference is played by causal anchors, variables that are known *a priori* to be causally upstream of others, and can be used to orient the direction of causality between other, relevant variables. A major application of this principle has been found in genetical genomics or systems genetics: genetic variations between individuals alter molecular and organismal phenotypes, but not vice versa, so these quantitative trait loci (QTL) can be used as causal anchors to determine the direction of causality between correlated traits from population-based data (Schadt *et al*., 2005; Chen *et al*., 2007; Rockman, 2008; Li *et al*., 2010). Such pairwise causal associations can then be assembled into causal gene networks to model how genetic variation at multiple loci collectively affect the status of molecular networks of genes, proteins, metabolites and downstream phenotypes (Schadt, 2009).

Interestingly, several experiments have recently shown that enhancer regions can be transcribed to form short, often bi-directional transcripts, called enhancers RNAs or eRNAs (Natoli and Andrau, 2012). eRNA expression is correlated with and crucially, *precedes* the expression of target genes (Arner *et al*., 2015). Though the functional role of eRNAs remains to be understood, the presence of eRNA from a regulatory region is an indicator of enhancer activity (Danko *et al*., 2015), and eRNA expression has been successfully used to predict transcription factor activity (Azofeifa *et al*., 2018). We therefore hypothesized that eRNA expression as a readout of enhancer activity could act as a causal anchor, opening new avenues to reconstruct causal gene regulatory networks.

To test this hypothesis, we developed novel statistical models and likelihood-ratio tests for using (continuous) eRNA expression data in causal inference, based on existing methods for discrete eQTL data, and implemented these in the Findr software (Wang and Michoel, 2017). We then applied this new method to CAGE data generated by the FANTOM5 project, a unique resource of enhancer and promoter expression across hundreds of human and mouse cell types (Forrest *et al*., 2014; Arner *et al*., 2015), and validated predictions using ChIP-seq and perturbation data. We noted that continuous eRNA expression values increased target prediction performance for some factors, while for other factors, a binarized presence or absence of the enhancer signal performed better. Leveraging this observation, we found that a data-driven approach to classify enhancer expression as either binary or continuous was sufficient to automatically select the best target prediction method, allowing parameter-free application of the method to organisms and cell types where validation data is not currently available.

## 2 Approach

We used enhancer expression as a causal anchor to infer causal gene interactions, within the Findr framework (Wang and Michoel, 2017). Findr provides accurate and efficient inference of gene regulations using eQTLs as causal anchors by accounting for hidden confounding factors and weak regulations. This is achieved by performing and combining five likelihood ratio tests (Figure 1B), each of which consists of a null (*ℋ*_null_) and an alternative (*ℋ*_alt_) hypothesis, to support or reject the causal model *E* → *A* → *B*, where E is an eQTL/enhancer in the regulatory region of gene *A*, and *B* is a putative target gene: primary linkage (*E* → *A*), secondary linkage (*E* → *B*), conditional independence (*E* → *B* only through *A*), *B*’s relevance (*E* → *B* or correlation between *A* and *B*), and excluding pleiotropy (partial correlation between *A* and *B* after conditioning on *E*). The log-likelihood ratios (LLRs) are computed for all possible targets of each gene, and then converted into *p*-values and posterior probabilities (Storey and Tibshirani, 2003) of the alternative hypothesis being true; see Wang and Michoel (2017) for details.

**Fig. 1.**
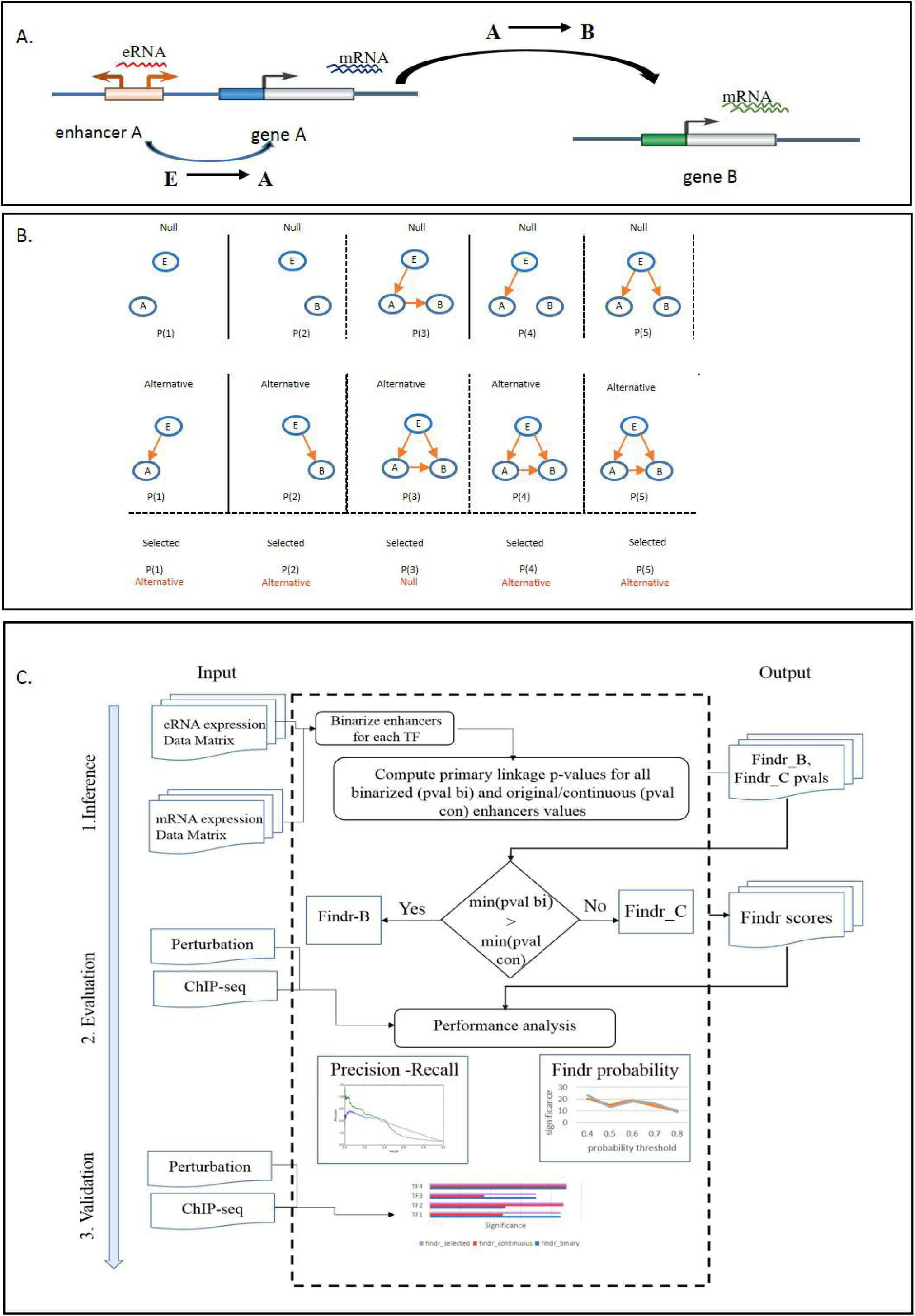
Overview of Findr framework. A. The schematic representation of causal gene regulatory network inference using enhancer activity as a causal anchor B. Five statistical tests used by Findr for causal inference C. work flow of the Findr-A framework.

We applied three treatments to enhancer expression data. First, we regarded enhancers as binary (on/off) variables, and after binarizing the data (see Methods), used the existing Findr to predict TF targets directly. This approach will be referred to as Findr-B (“binary”). Second, we adapted all five tests in Findr to use continuous instead of discrete causal anchor data, and used this method on (untransformed) eRNA data. This approach will be referred to as Findr-C (“continuous”). Third, to accommodate the co-existence of binary and continuous enhancers for different TFs within the same dataset, we developed an automatic adaptive method to independently treat each enhancer as binary or continuous, depending on the relative strength of the primary enhancer-TF linkage with either method. We call this approach Findr-A (“adaptive”).

## 3 Methods

### 3.1 Datasets

- We used Cap Analysis of Gene Expression (CAGE) data (TPM expression values) from the FANTOM5 Consortium for enhancer and transcription start sites (TSS) in mouse embryonic stem (ES) cells (36 experiments), macrophages (224 experiments) and erythroblastic leukemia (52 experiments) (Forrest *et al*., 2014; Arner *et al*., 2015). We also selected 1036 samples from all cell types and tissues in mouse.
- We use Chip-seq data from the Codex consortium (Sánchez-Castillo *et al*., 2015). Data available for 78 TFs in mouse ES cells, 12 TFs in macrophages and 17 TFs in erythroleukemia cells were used.
- For validation using Knock-out data, we have collected differentially expressed gene lists after perturbation of factors from published studies in mouse and gene lists after over-expression of factors in mouse ES cells from Xu *et al*. (2013).
- We obtained CAGE data (TPM expression values) from the FANTOM5 Consortium for enhancer and TSSs for human cell types and tissues (Forrest *et al*., 2014; Arner *et al*., 2015). We selected 360 samples from all cell types and tissues (1826 in total), by removing technical and biological replicates.

### 3.2 Data processing

- CAGE data was processed to clear for unannotated and non-expressed genes, and expression levels were log-transformed. Only enhancers expressed in more than one third of experiments were retained.
- For each TF, we selected the promoter with the highest median expression level as the promoter for that TF.
- For each TF, all enhancers within 50 kb of the TF promoter region were detected using the GenomicRanges package in Bioconductor Lawrence *et al*. (2013), and considered as candidate causal anchor enhancers for that TF.
- Enhancer data was binarized by setting all experiments with zero read count to 0 and all others to 1.
- For the ChIP-seq data, genes with a TF binding site within 1kb of their TSS were defined as targets for that TF.
- For the knockout and over-expression data, genes with differential expression *q*-value< 0.05 were defined as targets for the TF.

### 3.3 Likelihood ratio tests with continuous causal anchor data

Given a causal relation *E* → *A* → *B* to test, where *E* is a (continuous) enhancer for TF A and B is a putative target gene, with their expression data samples 1, …, *n* annotated in subscripts, we first convert each continuous variable into a standard normal distribution by rank. Each variable is modeled as a normal distribution with the mean linearly and additively dependent on its regulators in the five tests below (illustrated in Figure 1B).

1. **Primary linkage test**: The primary linkage test verifies that the enhancer *E* regulates the regulator gene *A*. Its null and alternative hypotheses are

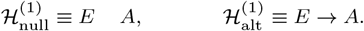 The log likelihood ratio (LLR) and its null distribution is identical with the correlation test in Wang and Michoel (2017). Therefore the LLR is

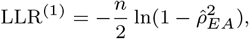

where

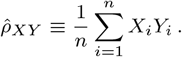 And its null distribution is

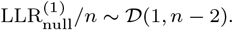 The probability density function (PDF) for 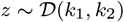 is defined as: for *z* > 0,

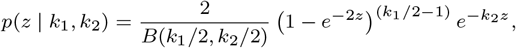

and for *z* ≤ 0, *p*(*z* | *k*_1_, *k*_2_) = 0, where *B*(*a,b*) is the Beta function.
2. **Secondary linkage test**: The secondary linkage test verifies that the enhancer *E* regulates the target gene *B*. The LLR and its null distribution are identical with those of the primary linkage test, except by replacing *A* with *B*.
3. **Conditional independence test**: The conditional independence test verifies that *E* and *B* become independent after conditioning on *A*, with its null and alternative hypotheses as:

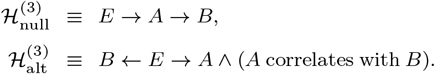 Correlated genes are modelled as having a multi-variant normal distribution whose mean linearly depends on their regulator gene. Therefore,

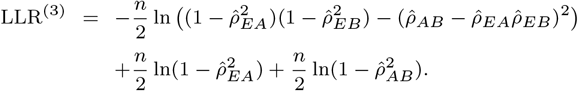 Following the same definition of the null data, its null distribution is

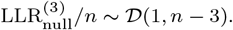
4. **Relevance test**: The relevance test verifies that *B* is regulated by either *E* or *A*. Its hypotheses are

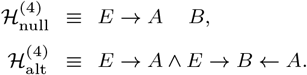 Similarly,

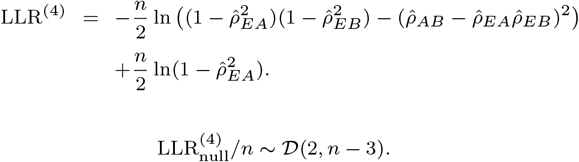
5. **Controlled test**: The controlled test verifies that *E* regulates *B* through *A*, partially or fully, with the hypotheses as

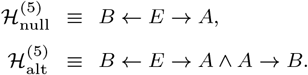 Its LLR is

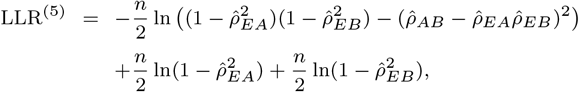

with the null distribution

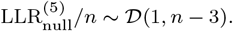 The LLR and its null distribution then allow to compute the p-values and the posterior probabilities of the null and alternative hypotheses separately for each subtest, as detailed in Wang and Michoel (2017).

### 3.4 Findr-B and Findr-C

In Wang and Michoel (2017), it was shown that a combined causal inference test performs best in terms of sensitivity and specificity for recovering true regulatory interactions, using both real and simulated test data. The combined test score is:

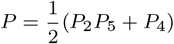

where *P_i_* is the posterior probability for subtest *i*.

The Findr-B method returns this combined *P*-value using the original Findr on binarized enhancer data. Findr-C does the same using the new tests on continuous enhancer data.

### 3.5 Adaptive method Findr-A

Given a set of TFs and for every TF, a set of candidate causal anchor enhancers, the adaptive Findr-A method performs the following, for each TF *A* (Figure 1C):

1. Compute the primary linkage test *p*-value for all candidate enhancers of *A*, both continuous and binarized.
2. Find the enhancer *E* with the lowest *p*-value overall.
3. If the lowest *p*-value occurred for a binarized enhancer, use Findr-B for TF A with *E* as its causal anchor, else use Findr-C.

### 3.6 Validation methods

For the purpose of evaluation, we calculated the Findr-B and Findr-C scores for all TF-gene combinations. Genes with scores exceeding a certain threshold were considered as predicted targets for each TF. Precision-Recall curves, calculated using the the “PRROC” package, and hypergeometric overlap *p*-values with respect to known targets from ChIP-seq and knockout data in the same cell type were used to compare the performance of the two methods.

## 4 Results

### 4.1 Causal inference from enhancer and transcript/gene expression CAGE data

To test our hypothesis that causal inference using enhancer expression as a causal anchor predicts true TF targets, we used CAGE data generated by the FANTOM consortium across hundreds of cell types and tissues in human and mouse (Forrest *et al*., 2014). We used bi-directional expression in nonpromoter regions as an indicator of likely enhancer activity in each CAGE sample (Andersson *et al*., 2014). The presence of enhancer expression (as a proxy for enhancer activity) in each sample is crucial for the ability to apply causal inference techniques. We first selected three mouse cell types (embryonic stem cells, macrophages and erythroblastic leukemia) for systematic characterization, as these had more than 20 samples per cell type, with diverse treatments or time series. Furthermore, ChIP-seq and TF knockout validation datasets were available for each of these cell types. We selected predicted enhancers (Andersson *et al*., 2014) within 50kb of each transcription factor in a cell type, resulting in 109 enhancers for 48 transcription factors in ES cells, 55 enhancers for 8 transcription factors in macrophages and 5 enhancers for 4 transcription factors in erythroleukemia, with an average of 3.8 enhancers per transcription factor across cell types.

We inferred causal transcription factor-target interactions for each transcription factor using the enhancer element most strongly linked to each TF in each cell type. We predicted targets for 48 transcription factors with two methods, one using continuous enhancer data (“Findr-C”), and one using discretized, binary (on/off) enhancer data (“Findr-B”) (see Methods). The Findr software outputs a score representing the putative probability of a causal interaction for each transcription factor-target pair (see Methods). For both methods, the targets with predicted probability of a causal interaction greater than 0.8 (see Methods) were validated using a compendium of ChIP-seq data (Sánchez-Castillo *et al*., 2015), containing 78 factors in ES cells, 12 factors in macrophages and 17 factors in Erythroblasts. Of these factors, 18 in ES cells, 7 in macrophages and 4 in Erythroblasts had enhancer expression in CAGE data. We noted that the suitability of Findr-B or Findr-C for causal inference was dependent on the factor, i.e. using continuous enhancer data performed better for some factors (Figure 2: Gata1, Fli1), while on/off data performed better for others (Figure 2: Myc,Klf2). Because the number of putative ChIP-seq targets for each factor varied widely across factors and cell types, from only 420 gene targets for JunD in Erythroblasts to over 12,000 gene targets for ESRRB in ES cells, the background precision levels differed highly between factors (Figure 2).

**Fig. 2.**
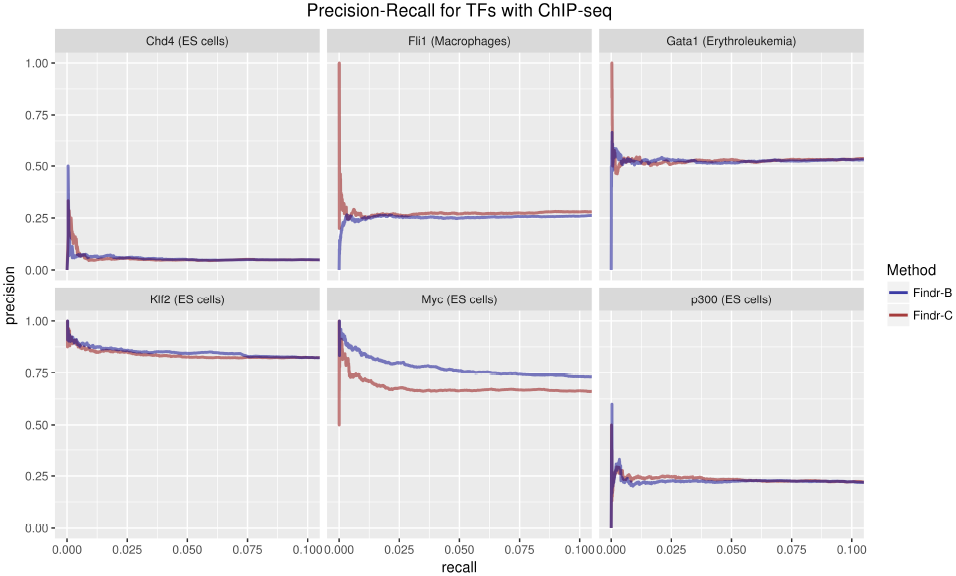
Recall-precision curves for target predictions by Findr-B and Findr-C using ChIP-seq and perturbation data

Transcription factor binding inferred using ChIP-seq data is thought to be mostly opportunistic and therefore might not to provide direct clues about the functional targets of the factor (Cusanovich *et al*., 2014). We therefore collected available perturbation data, specifically expression data after knock-out or knock-down (KO) of a factor, for 85 factors in ES cells and 11 factors in macrophages (see Methods). We also collected gene expression data after over-expression for 55 factors in mouse ES cells (Xu *et al*., 2013). Using differentially expressed gene lists as known targets, we evaluated the predictions of both methods. This confirmed the factor-specific suitability of either the Findr-B or Findr-C method (Figure 4).

We further tested whether these results were sensitive to the target probability prediction threshold. The enrichments of true positives were stable over a wide range around this threshold for both methods (Figure 3).

**Fig. 3.**
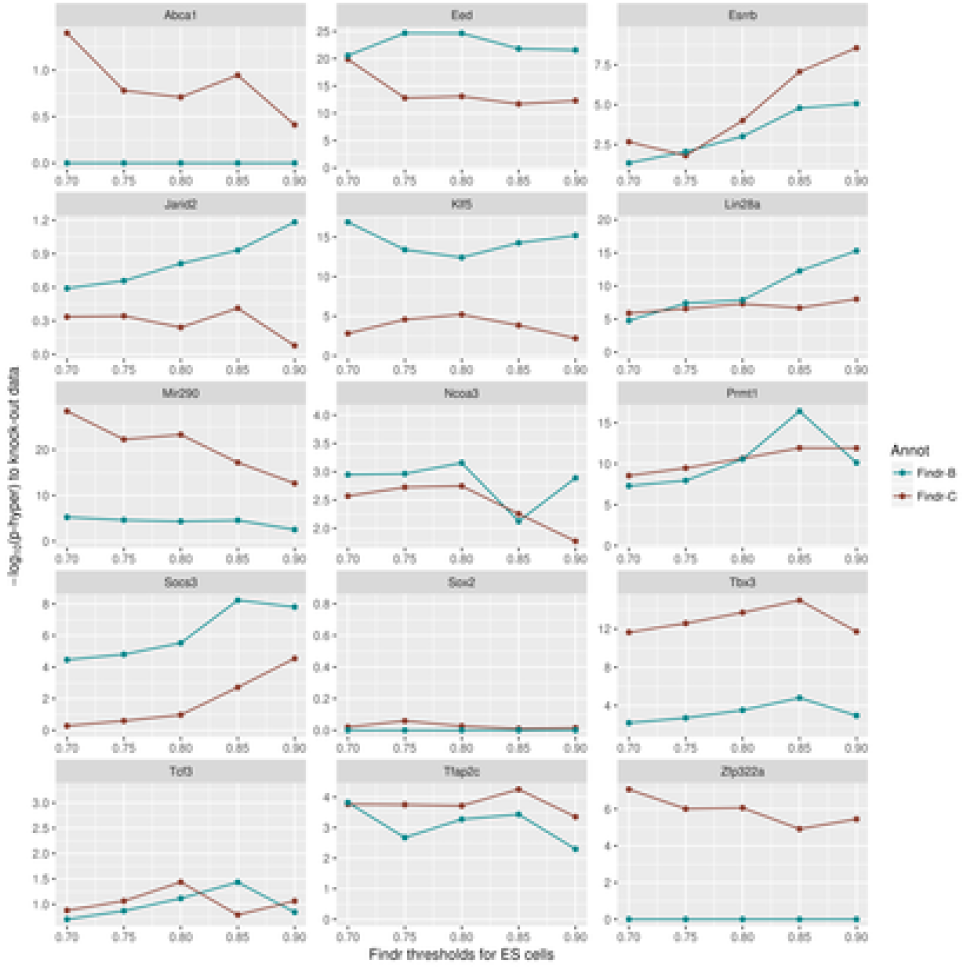
Robustness of Findr performance demonstrated by using different score thresholds

**Fig. 4.**
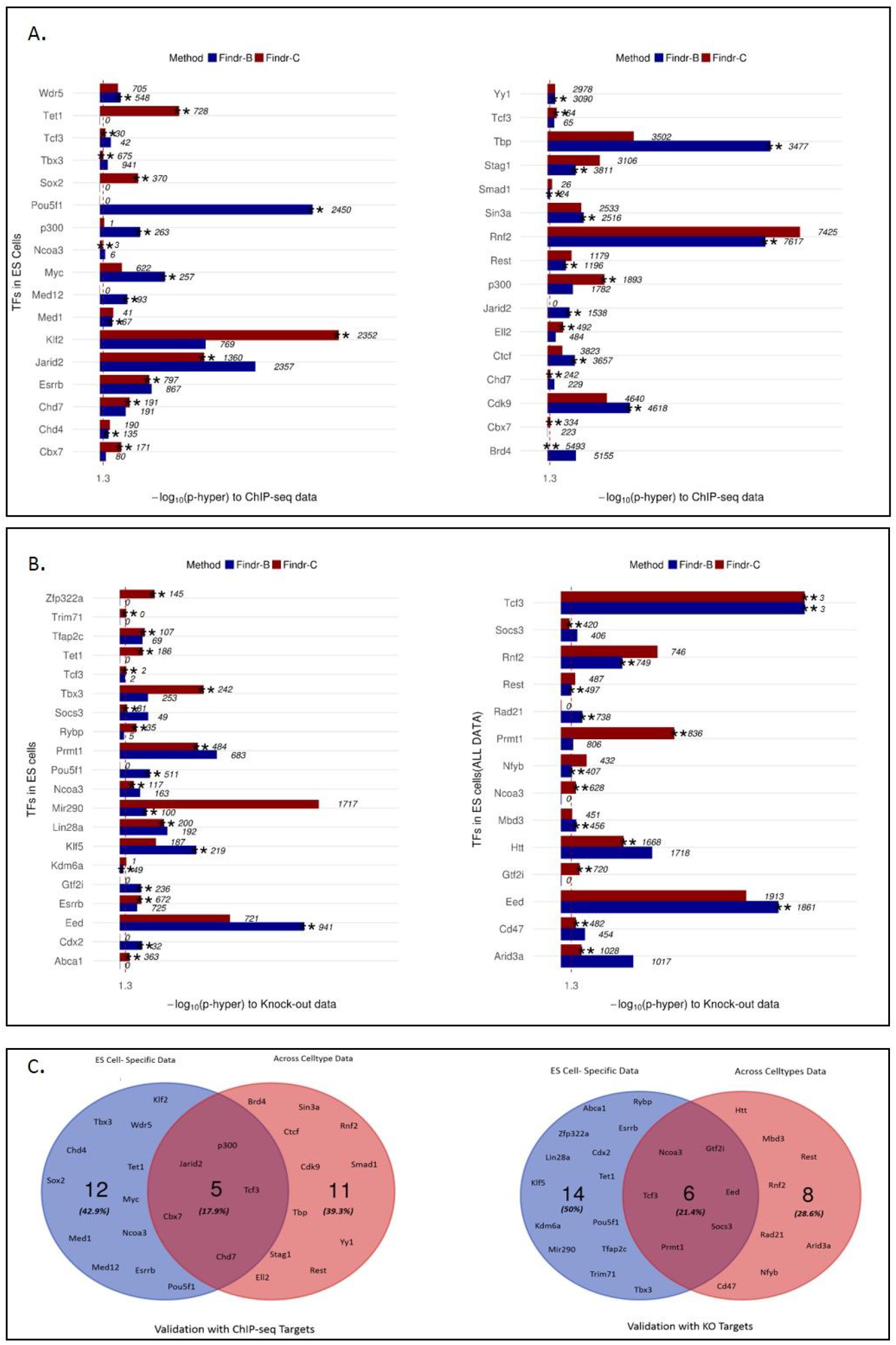
Comparison of Findr-A predictions using mouse ES cells and All cell types samples. A. bar plots representing enrichments for Findr-B, -C and -A predictions using ChIP-seq data as a validation dataset for ES cells (left) and all cell types (right). B. bar plots representing enrichments for Findr-B, -C and -A predictions using Knock-out data as a validation dataset for ES cells (left) and All cell types (right). C. overlap of factors between ES and All cell types using ChIP-seq (left) and knock-out (right) as validation datasets.

### 4.2 Development and validation of an adaptive model-selection approach for causal inference using discretized or continuous data

As the optimal prediction performance depended on a factor-specific choice between discretizing enhancer expression data or not, we investigated if this decision could be made in a data-driven, adaptive approach (henceforth called “Findr-adaptive” or “Findr-A”, see Figure 1C), in the absence of validation data. In short, Findr-A selects for each TF among all its candidate enhancers, both continuous and binary, the one with the strongest primary linkage to the TF’s expression, and then uses that enhancer and its corresponding method (Findr-B or Findr-C) to predict downstream targets for that TF (see Methods for details). This adapative approach was indeed able to select the best performing method for most of the factors (Figure 4A,B, Findr-A selection marked by a green star).

We further performed functional enrichment analysis of gene target sets predicted by Findr-A. 1119 targets of JunB in macrophages were enriched for ‘LPS signalling pathway’ (*p* < 10^−8^) and ‘RNA binding’ (*p* < 10^−8^). The macrophage CAGE samples indeed measured the response to LPS signalling. JunB is known to be a delayed response gene, attenuating transcriptional activity of immediate early genes and RNA binding proteins, specifically terminating translation of mRNAs induced by immediate early genes (Healy *et al*., 2013). 131 targets for Cbx7 in ES cells (Figure 3) were enriched for ‘regulation of transcription from RNA polymerase II promoter’ (*p* < 10^−10^), and included several developmental genes such as Hox family proteins. This enrichment was much stronger than for the ChIP bound targets of Cbx7 (*p* < 10^−5^). Cbx7 is a part of the PRC complex, which binds predominantly at bivalent chromatin at promoters of transcription regulators in ES cells (Mantsoki *et al*., 2015).

Traditional causal inference using causal anchors relies on a conditional independence test (Figure 1B, test 3), but the presence of common upstream regulatory factors results in low sensitivity for this test in applications to gene network inference (Wang and Michoel, 2017). To address this, both Findr-B and Findr-C (and hence also Findr-A) employ a combination of alternative tests (Figure 1B, test 2, 4 and 5, see Methods for details) that resulted in significantly improved prediction of TF targets from eQTL data (Wang and Michoel, 2017). This remained true for the CAGE data in all three mouse cell types (data not shown), and hence the combined test remains the recommended default.

### 4.3 Perturbations within and across cell types provide causal targets for a distinct set of transcription factors

We noted that most factors performed better using binary enhancer expression values and wondered if this might be due to the limited expression data available for each cell type. The Fantom5 CAGE data contains over 1000 samples across many more mouse cell types and tissues, henceforth called “all-data”. Findr-C indeed performed marginally better on all-data rather than cell type specific data. Importantly, all-data and cell type (ES) specific data resulted in causal targets for a distinct set of transcription factors. In particular, variation within a cell type was more informative for causal target predictions of cell type specific factors. For example, the targets of key pluripotency factors Sox2 and Esrrb (Dunn *et al*., 2014) were enriched in ES-data but not all-data (Figure 4A,B).

There were only five common factors with causal targets predicted using both all-data and ES-specific data that were validated by ChIP-seq data, and only six common factors validated by KO data (Figure 4C). Interestingly, the target genes predicted from ES-data and all-data for the same factor overlapped significantly. For example, 72% of Eed predicted targets using ES-data overlapped with Eed predicted targets using all-data.

### 4.4 Multiple enhancers of the same factor have a highly correlated expression

Mammalian genes are controlled by multiple enhancers. We investigated the stability of inference outcome under different choices of enhancers as causal anchors. For all factors for which Findr-A predicted targets in ES cells that overlapped significantly with perturbation data (Figure 4), we predicted additional target sets using other available enhancers, resulting in target sets for 76 enhancer-transcription factor pairs for 46 unique transcription factors.

The hierarchical clustering of transcription factor-enhancer pairs based on these target sets clustered mostly by transcription factors, indicating that the expression of multiple enhancers contains highly redundant information about the activity of the associated transcription factor (Figure 5). Reassuringly, factors known to form regulatory complexes, including Max and Mxi1 or Runx1 and Smad1, also clustered together, i.e. shared predicted causal targets (Figure 5).

**Fig. 5.**
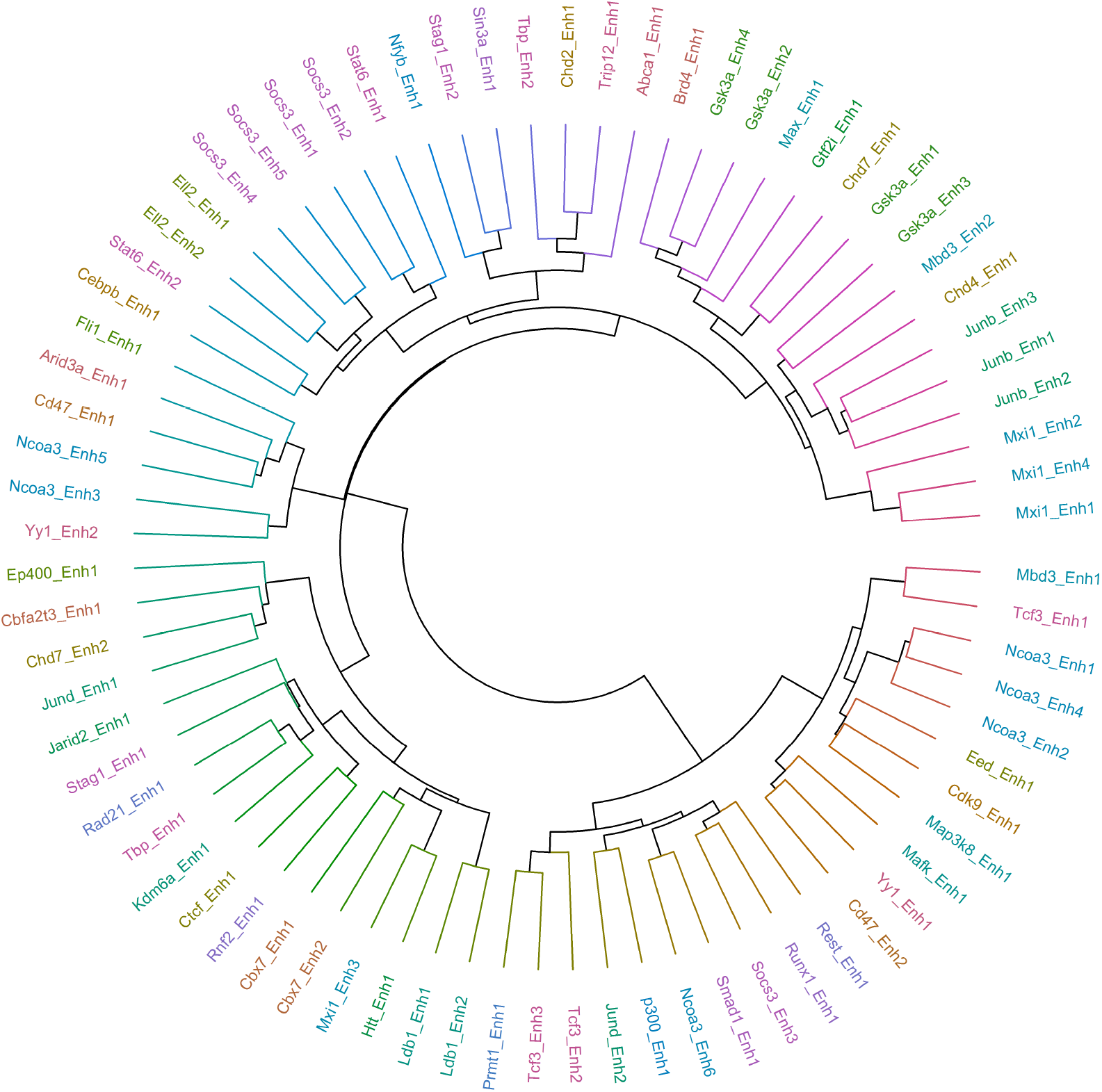
Circular plot of hierarchical clustering of overlap of targets predicted using multiple enhancers for the same factor in mouse ES cells

To investigate whether combining multiple enhancers was more informative for determining causal targets, we compared two integrative methods against the predictions of taking individual enhancers. Firstly, we used the median expression level of all the putative enhancers for each transcription factor as a ‘meta-enhancer’ in Findr-A. Secondly, we calculated the first principal component of the binary target prediction matrix for all enhancer of a TF in order to ‘average’ predictions. However, we did not observe any significant overall improvement in performance using either method.

### 4.5 Causal inference using CAGE expression data across human cell types

Finally, we inferred causal interactions between transcription regulators and targets using CAGE enhancer and TSS expression data in humans. Specifically, we inferred causal interactions for 20 transcription factors (with eRNA expression) using Findr-A (see Methods). We firstly validated the predicted interactions using a database of experimentally validated regulatory interactions in human (Han *et al*., 2018) and noted a statistically significant overlap between the predicted and experimentally validated gene sets (*p* < 10^−5^). Figure 6 represents the network of top 200 interactions for each factor. The factors involved in biologically related processes shared predicted causal targets. For example, two members of the SMAD family, SMAD3 and SMAD6, as well as BCOR and SIN3A involved in histone deacytelase activity showed a high overlap of predicted targets.

**Fig. 6.**
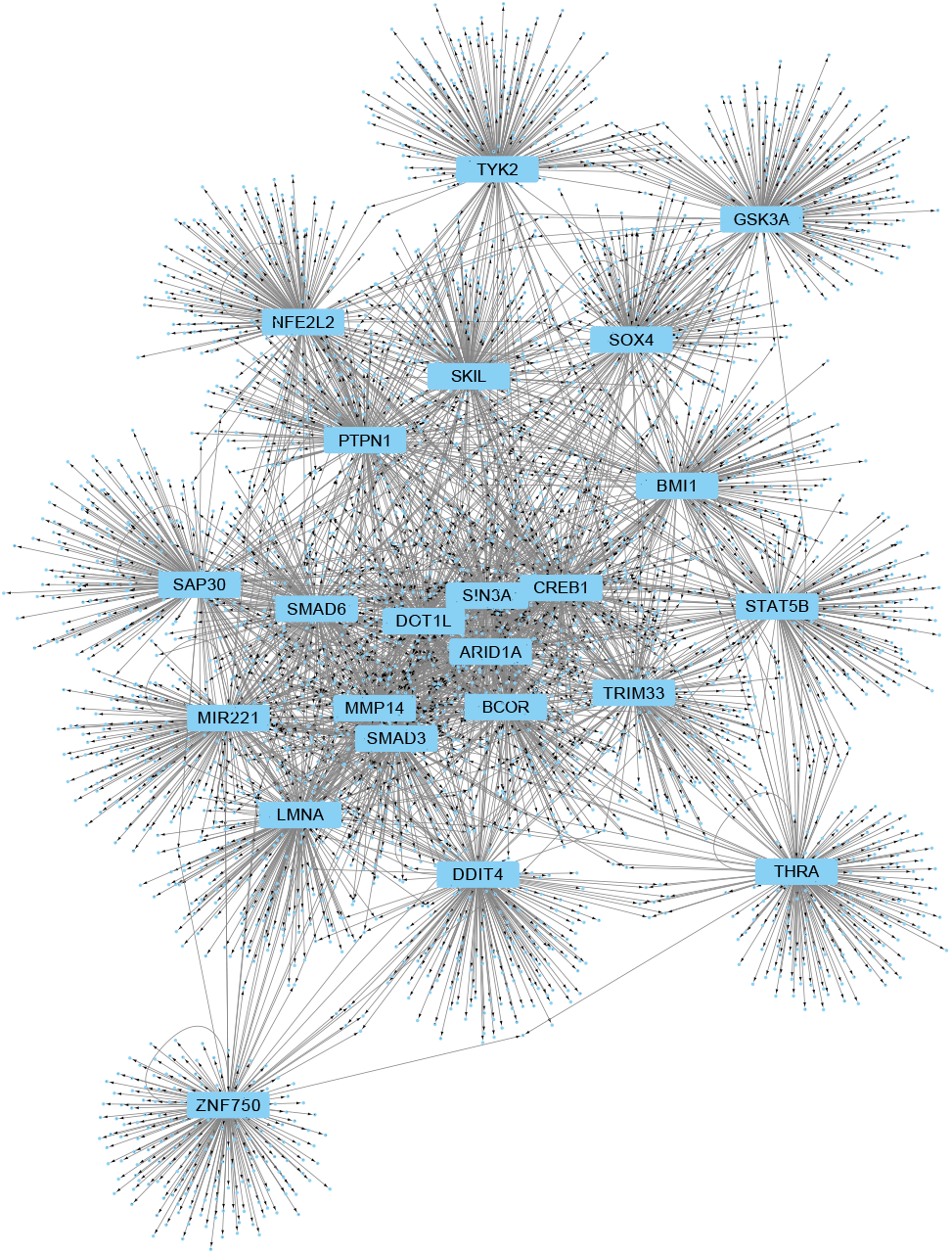
Network representation of transcription factor-targets predicted using Findr-A causal inference on FANTOM5 human dataset

## 5 Discussion & Conclusion

We explored the utility of eRNA expression as a causal anchor to predict transcription regulatory networks, by leveraging the observation that eRNAs mark the activity of regulatory regions. Previous studies support this notion, as eRNA expression has been shown to temporally precede the expression of its effector gene (Arner *et al*., 2015), and to correlate strongly with active regulatory regions across cell types (Danko *et al*., 2015). We therefore developed a novel statistical framework to infer causal gene networks (Findr-A), by extending the Findr software for causal inference using eQTL data (Wang and Michoel, 2017).

We demonstrated the applicability of Findr-A by predicting causal interactions from CAGE data generated through the FANTOM consortium, and validating them with ChIP-seq and perturbation data for three mouse cell types as well as on the entire FANTOM5 data. Notably, different factors were enriched for within cell type analysis as compared to across cell types. The causal regulatory network of cell type specific factors (e.g. Sox2, Esrrb in ES cells) could be inferred only using expression variation within a cell type and not across cell types. Due to the limited availability of validation data, a more comprehensive assessment was not possible.

The current approach can be extended in several aspects in the future. Firstly, Findr assumes equal (or no) relations between all sample pairs (Wang and Michoel, 2017), which hold for the majority of eQTL datasets. By accounting for heterogeneous sample relationships, such as biological and/or technical replicates, time series, or population structure, we may be able to reconstruct more accurate networks. Secondly, the assumption that eRNAs act as causal anchors is only approximately true, because their activity ultimately is regulated by other regulatory factors, i.e. the assumption that they are *a priori* causally upstream of correlated TF–gene pairs will not hold for all genes. Because eRNAs are temporally expressed before their direct target genes, we hypothesize that explicit modelling of gene expression dynamics in the Findr framework will allow to detect and correct for such feedback loops. Thirdly, eRNAs are expressed at relatively low levels, and therefore susceptible to noise, and a reliable eRNA signal was available for only a limited number of known transcription factors in mouse or human. Generating deep sequencing data for CAGE, utilizing GRO-seq or epigenetic or transcription factor ChIP-seq data to estimate enhancer activity could be possible ways to get around this.

In conclusion, we have demonstrated that enhancer activity can be used to infer causal gene regulatory networks. We foresee this approach to be of high value in the context of human medicine, by combining genetic, epigenetic and transcriptomic information across individuals to unravel causal disease networks.

## Funding

This work was funded by grants from the Biotechnology and Biological Sciences Research Council (BBSRC) [BB/P013732/1, BB/M020053/1].

## References

Andersson, R. et al. (2014). An atlas of active enhancers across human cell types and tissues. Nature, 507(7493), 455–461.

Arner, E. et al. (2015). Transcribed enhancers lead waves of coordinated transcription in transitioning mammalian cells. Science, 347(6225), 1010–1014.

Azofeifa, J. G. et al. (2018). Enhancer RNA profiling predicts transcription factor activity. Genome Res.

Chen, L. S. et al. (2007). Harnessing naturally randomized transcription to infer regulatory relationships among genes. Genome Biology, 8, R219.

Cusanovich, D. A. et al. (2014). The functional consequences of variation in transcription factor binding. PLoS Genet., 10(3), e1004226.

Danko, C. G. et al. (2015). Identification of active transcriptional regulatory elements from GRO-seq data. Nat. Methods, 12(5), 433–438.

Dunn, S. J. et al. (2014). Defining an essential transcription factor program for naïve pluripotency. Science, 344(6188), 1156–1160.

Forrest, A. et al. (2014). A promoter-level mammalian expression atlas. Nature, 507(7493), 462–+

Han, H. et al. (2018). TRRUST v2: an expanded reference database of human and mouse transcriptional regulatory interactions. Nucleic Acids Res., 46(D1), D380–D386.

He, B. et al. (2014). Global view of enhancer-promoter interactome in human cells. Proceedings of the National Academy of Sciences, 111(21), E2191–E2199.

Healy, S. et al. (2013). Immediate early response genes and cell transformation. Pharmacol. Ther., 137(1), 64–77.

Kundaje, A. et al. (2015). Integrative analysis of 111 reference human epigenomes. Nature, 518(7539), 317–330.

Lawrence, M. et al. (2013). Software for computing and annotating genomic ranges. PLoS Comput. Biol., 9(8), e1003118.

Li, Y. et al. (2010). Critical reasoning on causal inference in genome-wide linkage and association studies.

Mantsoki, A. et al. (2015). CpG island erosion, polycomb occupancy and sequence motif enrichment at bivalent promoters in mammalian embryonic stem cells. Sci Rep, 5, 16791.

Mifsud, B. et al. (2015). Mapping long-range promoter contacts in human cells with high-resolution capture Hi-C. Nature Genetics, 47(6), 598–606.

Natoli, G. and Andrau, J. C. (2012). Noncoding transcription at enhancers: general principles and functional models. Annu. Rev. Genet., 46, 1–19.

Pearl, J. (2009). Causality. Cambridge university press.

Rockman, M. V. (2008). Reverse engineering the genotype-phenotype map with natural genetic variation. Nature, 456(7223), 738–744.

Sánchez-Castillo, M. et al. (2015). CODEX: A next-generation sequencing experiment database for the haematopoietic and embryonic stem cell communities. Nucleic Acids Research, 43(D1), D1117–D1123.

Schadt, E. E. (2009). Molecular networks as sensors and drivers of common human diseases. Nature, 461, 218.

Schadt, E. E. et al. (2005). An integrative genomics approach to infer causal associations between gene expression and disease. Nature Genetics, 37(7), 710–717.

Storey, J. D. and Tibshirani, R. (2003). Statistical significance for genomewide studies. Proceedings of the National Academy of Sciences, 100(16), 9440–9445.

Thurman, R. E. et al. (2012). The accessible chromatin landscape of the human genome. Nature, 489(7414), 75–82.

Wang, L. and Michoel, T. (2017). Efficient and accurate causal inference with hidden confounders from genome-transcriptome variation data. PLOS Computational Biology, 13(8), e1005703.

Xu, H. et al. (2013). ESCAPE: database for integrating high-content published data collected from human and mouse embryonic stem cells. Database (Oxford), 2013, bat045.

